# Taxonomic profilers and their influence on metagenomic diversity analyses

**DOI:** 10.64898/2026.05.27.727884

**Authors:** Jonathan Rondeau-Leclaire, F. Guillaume Blanchet, Pierre-Étienne Jacques, Isabelle Laforest-Lapointe

## Abstract

Estimating taxonomic profiles is a central task in microbiome research. Several bioinformatic tools have been developed for this purpose, differing in algorithmic strategy, reference database flexibility, sensitivity parameters, and the type of abundance they estimate. As a result, taxonomic profiles carry an unwanted methodological signal whose driving characteristics remains understudied. While benchmarks have evaluated the performance of some of these tools, they rely on simulated data; little work has been done to compare them using real metagenomes in the presence of noise and uncharacterised diversity. Overall, the impact of taxonomic profiler choice and parameterisation on scientific conclusions remains poorly understood.

First, we provide a much-needed characterisation of four taxonomic profilers to help researchers better understand the available bioinformatic tools and inform their methodological choices. Then, we leverage 1,211 shotgun metagenomes from eight datasets to compare these taxonomic profilers across 13 methodological designs. Based on diversity indices, we found substantial variability in estimated taxonomic composition depending on methodological features such as reference database and algorithmic strategy. Alpha diversity analysis was substantially sensitive totool choice (particularly among *k*-mer-based tools) and reference database. Beta diversity showed sensitivity to both database and parameter choices, yet this variability barely affected statistical inference.

Our findings highlight the sensitivity of taxonomic diversity analyses to taxonomic profiling methodology and the importance for researchers to consider assessing the robustness of their results to choice of tool, parameter, and reference database. Crucially, differences in sample diversity across methodologies are symptomatic of differences in estimated taxonomic composition, which can affect any analysis based on taxonomic abundances. Overall, this study underscores the importance of tool selection and parametrisation, and of conducting sensitivity analyses to support robust and reliable scientific conclusions.

**AUTHOR SUMMARY:** Microbiome research relies on bioinformatic tools to determine which microbes are present in a sample and estimate their relative abundances, a process known as *taxonomic profiling*. Because this task requires comparing tens of millions of DNA sequences against thousands of microbial genomes, a wide range of computational strategies have been developed to make profiling accurate and efficient. As a result, many taxonomic profilers are now available, yet do not provide the same results. Although previous studies have evaluated their precision and sensitivity, less attention has been given to how the choice of profiler may influence the scientific conclusions drawn from microbiome data. Here, we selected four widely used taxonomic profilers to examine whether researchers would reach the same biological conclusions when comparing microbiomes across groups of samples, such as healthy and diseased individuals. We show that different profilers can lead to different biological conclusions and identify key tool characteristics that contribute to these discrepancies. Moreover, we provide a comparative overview of these four profilers, offering practical guidance for researchers seeking to choose the most appropriate tool for their study. These findings highlight the importance of software choice in microbiome research and support more transparent and reproducible data analysis practices.

## INTRODUCTION

The last decade saw the rise of metagenomics as a central approach in microbiome research. As sequencing costs decreased, the accessibility of metagenomics increased, with the total number of samples deposited in the NCBI Sequence Read Archive nearly quadrupling between 2020 and 2023 [1]. To process this growing, massive body of data, bioinformatic tools are being developed, published and updated regularly. As a result, researchers must make methodological choices at nearly every step of data processing, with the potential of affecting measurements and scientific conclusions. Here, we focus on algorithms designed to estimate microbiome community composition from shotgun metagenomes, often referred to as *taxonomic profilers*. Our aim is to help researchers understand the differences between popular tools and evaluate how bioinformatic parameters can affect the estimation of taxonomic composition diversity-based analyses.

Recent efforts from the field of metagenomics have provided recommendations for data processing [2–5] and classification [6–11], machine learning [12,13], statistical analyses [14–17], and general best practices [14,18–22]. Guidance based on systematic benchmarking have also emerged to help researchers choose or refine their bioinformatic methodology [23–26]. The Critical Assessment of Metagenome Interpretation (CAMI), a large scale effort to create a publicly available benchmarking framework for metagenomics, has standardised the evaluation of bioinformatic tools using simulated datasets [24,25]. CAMI, as well as other benchmarking studies [10,27,28], offers critical insights into accuracy and performance and highlight the context-dependency of what constitutes a “better tool”, underscoring that there is no one-size-fits-all methodology in metagenomics [29]. Overall, what constitutes the “right” tool depends on multiple factors such as a sample’s origin, complexity, phylogenetic diversity, and sequencing depth. Although many tools perform well in estimating taxonomic profiles, choosing the right one and using it adequately remains a challenge for researchers.

Benchmarking metagenome taxonomic profilers requires a known ground truth, often taking the form of simulated metagenomes. With over 95% of prokaryotic species being genomically uncharacterised[30], simulated metagenomes are not likely to well represent real microbiomes, especially those of understudied environments or hosts. Moreover, benchmarking tools against one another implicitly assumes that any two “good enough” tools should produce converging outputs. Yet for taxonomic profilers, it is not well established whether all tools *should* yield the same result, and therefore whether they are strictly comparable. Importantly, every method has its own biases and limitations, and some authors have argued against methodological standardisation [29,31], favouring instead more transparent reporting [32]. To better select and justify taxonomic profiling methodology, researchers thus need a clear understanding of what is estimated, the methodological limits, and the extent to which these tools can be used interchangeably. In this paper, we qualitatively compare the characteristics of four taxonomic profiling tools. Then, we quantitatively assess how user choices influence sample diversity estimates and statistical comparisons using real metagenomes.

### TAXONOMIC PROFILING TOOLS

Several taxonomic profilers for shotgun metagenomics have been published over the last decade. These tools compare reads to a set of reference sequences, producing either *taxonomic* or *sequence* relative abundances [33]. Tools that use a DNA-to-markers (D2M) approach align metagenome reads to their own collection of genes and estimate relative *taxonomic* abundances by normalising the number of aligned reads by gene length. Two popular tools based on that approach are MetaPhlAn [34] (Metagenomic Phylogenetic Analysis) and mOTUs [35] (Marker gene-based Operational Taxonomic Unit). Alternatively, Kraken2 [36] and Sourmash [37,38] use a DNA-to-DNA (D2D) approach, meaning they rely on matching the *k*-mers of metagenome reads to a collection of genomes [7], and thus estimate relative *sequence* abundances. These are not normalised by genome length, as this information is unknown for incomplete metagenome-assembled genomes (MAGs) and becomes ambiguous at higher taxonomic ranks (which Kraken assigns in its algorithm, as opposed to Sourmash). Therefore, sequence abundances are biased by true genome length, as longer genomes will produce more reads than shorter genomes [33]. Despite this, tools from both the D2M and D2D approaches are widely used in microbiome research.

Importantly, it must be highlighted that distinct abundance types produce different taxonomic profiles that are not equivalent and cannot be interconverted [39,40]. This recently exposed nuance [33] between the D2M and D2D tools can lead to incorrect comparisons, as both approaches cannot yield the same abundance profiles. However, the causes of this divergence can go beyond the type of abundance estimated. These tools can also differ in their algorithmic strategy, database flexibility, reference taxonomy, and strategy for detecting unknown species. **Table 1** summarises the characteristics of these tools.

**Table 1.** Characteristics of selected taxonomic profiling tools.

|  | **mOTUs4** | **MetaPhlAn** | **Kraken2** | **Sourmash** |
| --- | --- | --- | --- | --- |
| **Approach & abundance type** | D2Ms,<br>taxonomic abundances |  | D2D,<br>sequence abundances |  |
| **Reference database** | 10 universal marker genes | >5 million clade-specific marker genes | Genomes (including MAGs) |  |
| **Reference database flexibility** | Version + user-added genomes | Version only | Full (requires full re-indexing) | Full (requires appending only) |
| **Taxonomy** | NCBI<sup>1</sup> |  | Follows reference database (default: NCBI) | Follows reference database |
| **Unknown species** | meta-mOTUs & ext-mOTUs<sup>2</sup> | uSGBs | User-added MAGs |  |

| Sample<br>identifiability<br>metric | Estimated proportion<br>of unknown taxa |  | % classified reads | % metagenome<br>containment |
| --- | --- | --- | --- | --- |
| Algorithmic<br>strategy | Read alignments<br>(BWA-MEM) | Read alignments<br>(Bowtie2) | *k*-mer matching:<br>Lowest common<br>ancestor | *k*-mer matching:<br>Minimum<br>metagenome cover |

<sup>1</sup> tool for mappings to GTDB provided
<sup>2</sup> mOTUs has an extension tool allowing users to find a label mOTUs in MAGs they provide
MAG: Metagenome-Assembled Genome; SGBs: Species-Level Genome Bins; uSGBs: unknown SGBs.

### Unknown species detection strategy and database flexibility

The D2M tools evaluated here can detect many species lacking a reference genome or taxonomy. Both MetaPhlAn and mOTUs include in their reference database genes from uncharacterised species, extracted from public metagenomes. However, these tools employ fundamentally different strategies to find unknown species. MetaPhlAn takes a “genome first” approach, clustering over a million reference genomes and MAGs into known species-level genome bins (kSGBs, containing at least one reference genome) and unknown SGBs (uSGB, containing only MAGs). These SGBs are then used to identify over 5 million clade-specific marker genes to create a reference database. MetaPhlAn aligns metagenomic reads to these genes to determine sample taxonomic composition. Conversely, mOTUs takes a “marker first” approach, mining publicly available genomes and metagenomes for 10 universal phylogenetic marker genes. Markers found in reference genomes inherit their taxonomy directly and form ref-mOTUs, while markers found in metagenome assemblies and MAGs are clustered into species-level units (meta-mOTUs and ext-mOTUs, respectively) based on their similarity. These new units are then taxonomically labelled at higher rank based on their similarity to the closest known species, yielding a reference database of genes to which mOTUs aligns metagenomic reads.

In comparison, D2D tools such as Kraken and Sourmash do not have a built-in strategy for unknown species detection. However, since their database can be any set of taxonomically labelled genomes (often RefSeq or GTDB databases), users can assemble MAGs from the metagenomes whose composition they wish to estimate and build a custom reference database (or complement an existing one). This feature is especially useful for studying microbial communities whose members are poorly represented in public databases. Sourmash can append genomes to an existing index, whereas Kraken requires the entire database index to be rebuild, a more resource-hungry process. Of note, as of version 4, mOTUs [41] can now find marker genes in user-provided genomes, to add them to the reference database using an extension program, though this feature is not evaluated here.

### Algorithmic approaches to k-mer matching

The D2D tools evaluated here differ mainly in terms of algorithmic strategy. Both Kraken and Sourmash rely on matching k-mers (each “word” of length *k* in a read) exactly between reads and reference genomes. Essentially, reads are assigned to sequences with which they share the most k-mers; however, each tool uses different heuristics to accomplish this task.

Briefly, Kraken2 classifies a read using the genome that has the highest number of exact *k*-mer matches with that read. When a read’s *k*-mers matches two genomes equally well, Kraken2 assigns it the lowest taxonomic rank [42] these genomes have in common. Abundances are based on the number of *k*-mers per taxonomic rank and are usually re-estimated using Bracken, a tool that redistributes reads from higher taxonomic ranks down to the desired level (e.g., Species) based on Bayes’ theorem. It is generally recommended to use Bracken when estimating taxonomic compositions with Kraken, because it helps increase the taxonomic classification rate at the species level. Because it classifies each read independently, Kraken2 defines the confidence required for a read to be labelled (the *--confidence* parameter). Importantly, even if a reference genome is only matched by one read from a metagenome, that taxon will be reported as part of its composition (though at extremely low abundance).

Sourmash takes a different approach: it finds the smallest set of reference genomes that contain all the identifiable *k*-mers of a metagenome [38]. To do this, it finds the genome covering the largest amount of *k*-mers, removes these *k*-mers from the query (the metagenome), then repeats this process iteratively. Sourmash estimates abundances based on the number of k-mers uniquely matched to all detected genomes and, contrary to Kraken, will not report a taxon matched by very few reads; its *--threshold_bp* parameter controls the minimum size for a *k*-mer set to be retained. Currently, there is no consensus on which parameter values are optimal across analytical contexts when comparing Kraken and Sourmash, although great efforts were made recently to benchmark Kraken’s *--confidence* parameter [39]. In both cases, default settings may not be optimal and their modification requires careful consideration of the trade-offs between sensitivity and specificity [27], an assessment delegated to the user.

### Reference database taxonomy

Two taxonomies dominate prokaryotic classification in metagenomics: NCBI and GTDB. Whereas NCBI taxonomy integrates historical nomenclature alongside phylogenetic data, GTDB is exclusively based on phylogenomic relationships and is designed to ensure monophyly across clades. However, GTDB does not always follow the *The International Code of Nomenclature for Prokaryotes* [43], which can be problematic in clinical or epidemiological contexts, depending on research objectives. Consequently, a metagenome’s observed diversity can easily be influenced by the taxonomy of the reference database. For example, if *Shigella* taxa are present in the sample, the relative abundance of *E. coli* will appear higher with GTDB than with NCBI as GTDB considers *Shigella* to be *E. coli* strains.

As they simply rely on a set of reference genomes, D2D tools allow the user to choose the reference taxonomy by selecting a RefSeq database (NCBI taxonomy) or a GTDB database. D2M tools, on the other hand, are restricted to databases specifically built by their authors, which thus constrains the taxonomy. It is therefore likely that genus or species diversity estimations will vary depending on which tool is used, and hypotheses based on diversity indices may reveal different conclusions accordingly. Of note, mOTUs provides a mapping file between NCBI and GTDB taxonomies, allowing the user to convert the output.

Overall, these differences in tool characteristics and algorithmic flexibility highlight the need to assess the extent to which taxonomic profilers diverge in their estimation of a metagenome’s taxonomic composition and identify which methodological features, beyond abundance type, contribute to these divergences. This knowledge will help guide methodological choices through a clearer understanding of the influence of tool characteristics on downstream analyses.

In this work, we first evaluate the sensitivity of intra-sample (alpha) and inter-sample (beta) diversity indices across four taxonomic profilers and methodological choices using real metagenomes. We then illustrate how such variability can influence the outcomes of hypothesis testing, before discussing which tool characteristics likely drive this sensitivity and proposing best practices to improve comparability, transparency and robustness in metagenomic research.

## METHODS

We analysed eight datasets comprising a total of 1,211 metagenomes (**Table 2**). Each contains two discriminating groups (e.g., sick vs. healthy) as defined by the original study authors. Sequences were quality-controlled and trimmed using Kneaddata [44] with parameters *SLIDINGWINDOW:4:20* and *MINLEN:50*. Human-associated datasets were decontaminated by removing human reads using the GRCh38 reference genome and Bowtie2 *--very-sensitive-local* mode. Environmental datasets were not decontaminated.

**Table 2.**
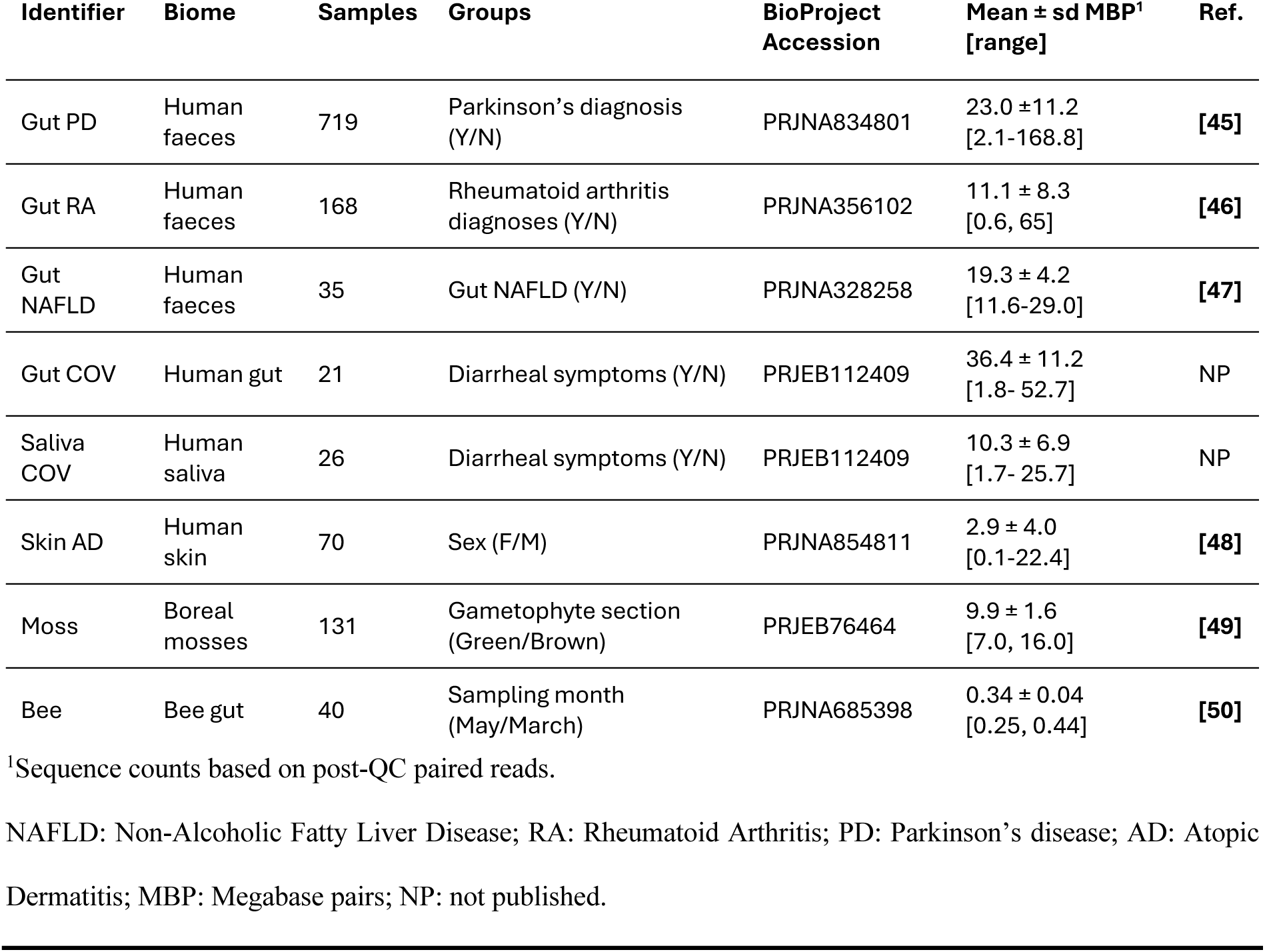
Description of datasets used.

### Taxonomic profiling tools

Four tools were compared: mOTUs v4.0.4 [41], MetaPhlAn v4.0.4 [34], Sourmash v4.8.11 [37,38,51] (the *gather* algorithm) and Kraken v2.1.2 [36]. A total of thirteen methodologies were tested, each combining one of these tools with variations in reference databases or tool-specific parameters (**Table 3**). Kraken was evaluated across a range of plausible values for the *--confidence* parameter.

**Table 3.**
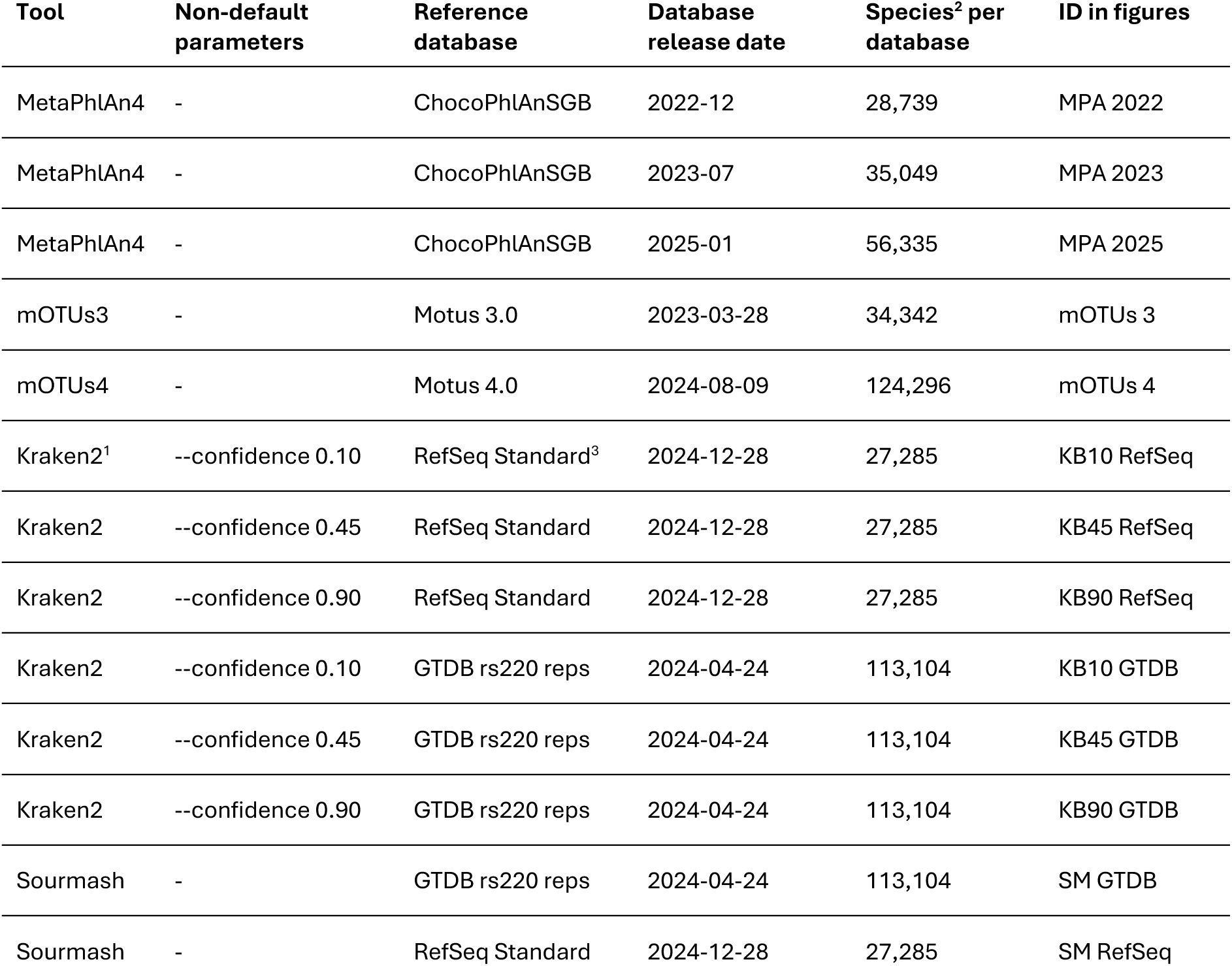
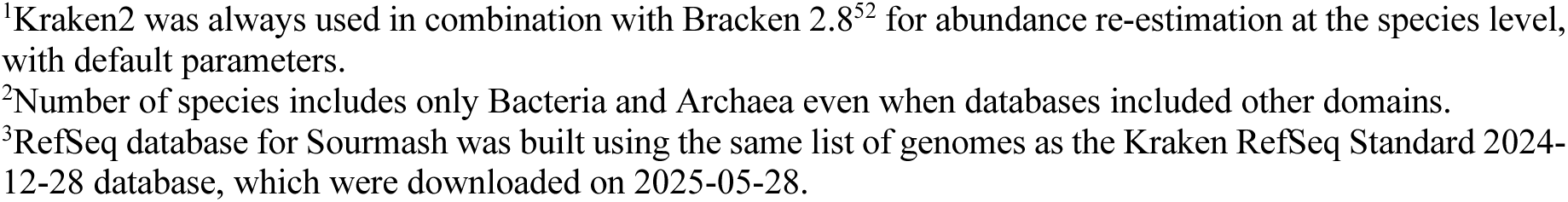
Taxonomic profiling methodologies evaluated in this study.

| Tool | Non-default parameters | Reference database | Database release date | Species<sup>2</sup> per database | ID in figures |
| --- | --- | --- | --- | --- | --- |
| MetaPhlAn4 | - | ChocoPhlAnSGB | 2022-12 | 28,739 | MPA 2022 |
| MetaPhlAn4 | - | ChocoPhlAnSGB | 2023-07 | 35,049 | MPA 2023 |
| MetaPhlAn4 | - | ChocoPhlAnSGB | 2025-01 | 56,335 | MPA 2025 |
| mOTUs3 | - | Motus 3.0 | 2023-03-28 | 34,342 | mOTUs 3 |
| mOTUs4 | - | Motus 4.0 | 2024-08-09 | 124,296 | mOTUs 4 |
| Kraken2<sup>1</sup> | --confidence 0.10 | RefSeq Standard<sup>3</sup> | 2024-12-28 | 27,285 | KB10 RefSeq |
| Kraken2 | --confidence 0.45 | RefSeq Standard | 2024-12-28 | 27,285 | KB45 RefSeq |
| Kraken2 | --confidence 0.90 | RefSeq Standard | 2024-12-28 | 27,285 | KB90 RefSeq |
| Kraken2 | --confidence 0.10 | GTDB rs220 reps | 2024-04-24 | 113,104 | KB10 GTDB |
| Kraken2 | --confidence 0.45 | GTDB rs220 reps | 2024-04-24 | 113,104 | KB45 GTDB |
| Kraken2 | --confidence 0.90 | GTDB rs220 reps | 2024-04-24 | 113,104 | KB90 GTDB |
| Sourmash | - | GTDB rs220 reps | 2024-04-24 | 113,104 | SM GTDB |
| Sourmash | - | RefSeq Standard | 2024-12-28 | 27,285 | SM RefSeq |

For Kraken, we used the publicly available GTDB rs220 species representative database and the 2024-12-28 RefSeq Standard database (available at https://benlangmead.github.io/aws-indexes/k2). For Sourmash, the GTDB rs220 index published by authors was used (available at https://sourmash.readthedocs.io/en/latest/databases-md/gtdb220.html), and a custom index was built using the same RefSeq Standard accessions found in the Kraken database. Sourmash databases were built with a *k*-mer size of 31 and a scaling factor of 1000. Furthermore, to increase comparability across methodologies, only bacterial and archaeal taxa were kept, the only domains represented in the GTDB database.

To control for uneven sequencing depth, abundance tables were rarefied to the smallest sample size of each dataset using the *rarefy_even_depth()* function from the *phyloseq* [52] package, in line with recent recommendations [14]. The tables produced by the MetaPhlAn methodologies were not rarefied, because they are already normalised as proportions (total sum scaling).

### Alpha diversity

Four alpha diversity metrics were calculated: richness [53], Shannon entropy [54], and the inverse of the Simpson index [55]. All three are widely used owing to their early adoption in microbial ecology studies. In addition, the Tail statistic [56], developed specifically for microbiome research was also used. Richness represents the number of detected species, which can be highly sensitive to sequencing depth and taxonomic profiling tool parameters. In the context of microbial compositions, the Shannon index quantifies the uncertainty in predicting the taxon identity of a randomly drawn sequence, with higher values reflecting greater diversity. The Simpson index is the probability that two randomly drawn sequences belong to the same taxa, with lower values indicating greater diversity. For ease of interpretation, we use the inverse Simpson index, representing the effective number of equally abundant taxa that would produce the same diversity. Finally, the Tail statistic is the weighted standard deviation of taxon ranks, with the distance of each rank measured to the most abundant taxon and weighted by its relative abundance. This statistic was developed to better reflect low-abundance taxa in a diversity index.

MetaPhlAn and mOTUs cannot interchange their reference database. Thus, diversity estimates can be affected simply by the number of available taxa in each database, and the impact of tool choice cannot be de-confounded from the impact of database choice within the D2M approach. Kraken2 and Sourmash, on the other hand, can use the same reference database. Even then, diversity estimates will be influenced by Kraken2’s *--confidence* threshold or Sourmash’s *--threshold-bp*, which have no single “optimal” value and directly affect the number of reported species. Therefore, to account for the large differences in representative species across reference databases and for the unmatched tool-specific detection sensitivity parameters in Kraken and Sourmash, diversity values were centred and scaled within methodology across all datasets, using the median and median absolute deviation (MAD) for robust scaling [57].

For each pair of comparable tools (MetaPhlAn vs. mOTUs, and Kraken2 vs. Sourmash using either GTDB or RefSeq as a reference database), the difference in scaled diversity was computed for each sample. The variance of these distributions was used as a measure of methodological agreement: a wider spread indicates that diversity estimates are more strongly influenced by the choice of tool, even after removing approach-specific effects. Differences in these variances were assessed statistically using the Pitman-Morgan test [58,59] for paired samples using the PairedData package [60] (v1.1.1).

To assess the impact of methodology on scientific conclusions based on alpha diversity comparison, for each index, a Mann-Whitney [61] test was performed for each methodology-dataset combination, evaluating the significance and effect size of differences in diversity between two groups of samples as defined by the dataset metadata (**Table 3**). Effect sizes are expressed as rank-biserial correlations [62].

### Beta diversity

Beta diversity is used to evaluate whether microbiota compositions differ more between groups of samples than within them. To that end, distance or dissimilarity indices can be used, which quantify the compositional similarity between every pair of samples. Here, we focus on the Bray-Curtis index because, in addition to being widely used, it accounts for relative abundances, which is something taxonomic profilers try to estimate. Alternative indices have also been developed to account for phylogenetic distances between taxa[63] or data compositionality [64,65], but their comparison is beyond the scope of this paper.

For each methodology-dataset combination, a between-sample Bray-Curtis dissimilarity matrix was computed. Ordinations were performed on each of these matrices using Principal coordinates analysis (PCoA, sometimes also referred to as metric multidimensional scaling [66]) using the *cmdscale* function from the stats packages in R. Separately for each dataset, Procrustes correlations [67] were computed between the ordinations of each pair of methodologies using the *protest* function of the vegan package [68] (v2.7-3). For each of these methodology pairs, we report the median correlation ± MAD. Methodologies were clustered hierarchically based on the median correlation, using the *hclust* function from the stats package [69] (v4.5.2). Finally, the eigenvectors of each Procrustes matrix were extracted to visually assess the clustering of methodologies for each dataset.

To assess the impact of methodology on scientific conclusions based on beta diversity comparison, we performed a permutational analysis of variance [70] (PERMANOVA) using the *adonis2* function of the vegan package on Bray-Curtis dissimilarities for each dataset-methodology combination using the model *Dissimilarity Matrix ∼ Group*. As for the Mann-Whitney test, P-values were not adjusted and are shown alongside the proportion of variance explained, quantified by a coefficient of determination (R^2^), which is the ratio of the between groups and total sum of squares.

## RESULTS

### Alpha diversity indices across methodologies

Alpha diversity for any given sample varied greatly, even between tools of the same approach or when controlling for reference database. With D2M tools, using the most recent databases, mOTUs4 produced higher Shannon (**Figure 1A**), Inverse Simpson (**Figure S2A**), and Tail (**Figure S3A**) diversities than MetaPhlAn4 for 95.5%, 89.0%, and 95.0% of samples, respectively. Conversely, 91.4% of samples had higher richness (**Figure S4A**) with MetaPhlAn4.

**Figure 1.**
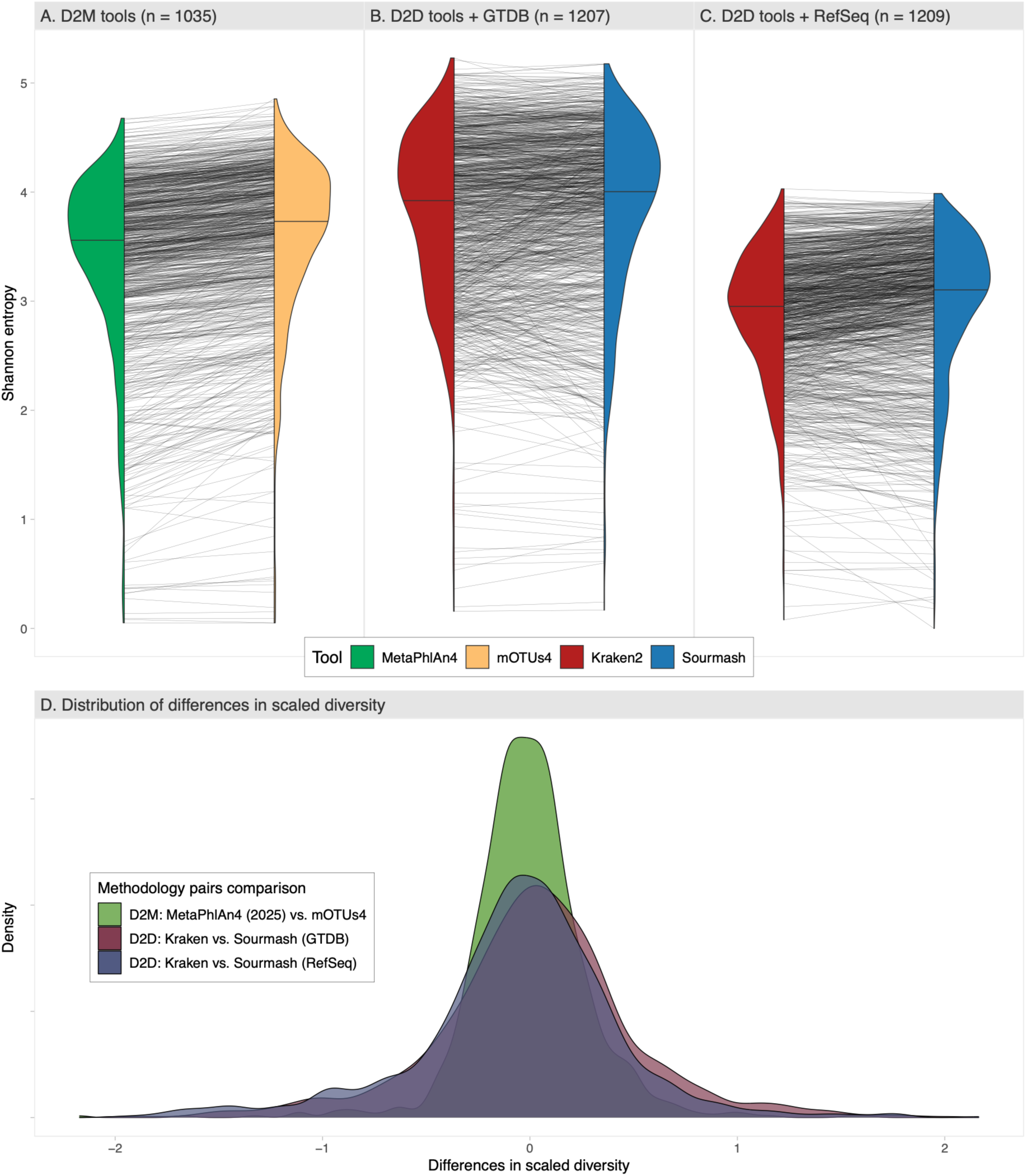
Changes in estimated Shannon diversity between pairs of methodologies having common characteristics over eight datasets. A-C) Lines connect a sample between methodologies. Half-violin plots show the density of samples. Out of all 1,211 samples used in this study, only those whose composition was successfully estimated by both tools in a pair are presented (n indicated in facet title). Kraken2’s confidence threshold was set to 0.45; D2M methods use the latest reference databases (**Table 3**). D) Differences in Shannon diversity scaled by methodology. D2M: DNA-to-markers; D2D: DNA-to-DNA.

When comparing D2D tools using the same reference database, every sample exhibited higher richness when taxonomic composition was estimated by Kraken at a confidence threshold of 0.45, compared to Sourmash (GTDB: 1167 ± 625.7 species, RefSeq: 192 ± 48.69 species; **Figure S4B,C**). Even at a stringent Kraken *--confidence* of 0.90, richness remained higher when evaluated by Kraken for 79.2% (GTDB) and 91% (RefSeq) of samples. When comparing reference databases using the same D2D tool, GTDB yielded substantially higher richness (Kraken 1213 ± 642 for 95% of samples; Sourmash 200 ± 141.9 for 99.9% of samples), which is expected, given that the GTDB 220 database contains over four times more species than the RefSeq database (**Table 3**) used in this study.

The differences between Kraken and Sourmash were much less consistent for indices based on relative abundances (Shannon **Figure 1B,C**; Inverse Simpson **Figure S2B,C**; and Tail **Figure S3B,C**). At first glance, this suggested tool-specific biases in diversity estimation. However, lowering Kraken’s *--confidence* parameter produces higher diversity values than Sourmash for an increasing proportion of samples across all indices. In fact, even when using a Kraken confidence level that generates a diversity distribution similar to Sourmash’s *(--confidence* 0.45 was a good approximation for Shannon), sample diversities still varied substantially in both directions depending on the chosen tool (**Figure 1B,C**).

### Sensitivity to approach and reference database

To assess the sensitivity of diversity indices to methodological choices while accounting for tool-specific algorithmic sensitivity parameters or difference in reference databases, we compared the distribution of same-sample differences in centred and scaled diversity measurements between pairs of methodologies (<u>see *Methods*</u>). Shannon and Inverse Simpson indices showed the largest differences in variance of within-sample differences when comparing D2M pairs to D2D pairs (**Figure 1D**; **Figure S2**; Paired Pitman-Morgan tests all p<0.001 **Table S1**). Between D2D tools, three indices showed higher variance when using RefSeq as a reference compared to GTDB (richness and Tail p<0.001; Simpson p=0.016) but not Shannon (p=0.234). Based on variance, alpha diversity indices showed the lowest sensitivity to choice of tool within the D2M approach, and the highest with tools of the D2D approach used with the RefSeq reference database.

### Beta diversity ordinations across methodologies

To assess how methodology influences beta diversity estimates, we compared the principal coordinates of within-dataset Bray-Curtis dissimilarity matrices between all methodology pairs using Procrustes correlations. A Procrustes analysis compares two ordination results by matching the shape of both ordinations in multivariate space using rotation, reflection and scaling, thus quantifying their correlation.

Methodologies using D2M tools or D2D tools with GTDB, with the exception of KB90 GTDB, showed median correlations between 0.92 and 1.00 (**Figure 2A**) with very low median absolute deviations, between 0.00 and 0.03 (**Figure S1**). Reassuringly, Procrustes correlations between D2M methodologies differing only by their database *version* were >0.98 (mOTUs 3 vs. 4 and MetaPhlAn 2023 vs. 2025), indicating near-perfect ordination match. Correlations between Sourmash and Kraken methodologies were highest when using GTDB (from 0.88 to 0.99), as opposed to RefSeq (from 0.82 to 0.93). Across all methodologies, KB90 RefSeq disagreed the most, with Procrustes correlations to all non-RefSeq methodologies ranging from 0.68 ± 0.07 (KB90 GTDB) to 0.75 ± 0.05 (KB10 GTDB). **Figure 2B** illustrates a PCoA displaying both tighter clustering and greater variance explained by the first two axes when RefSeq is used than when GTDB is used with Kraken on the Gut RA dataset.

**Figure 2.**
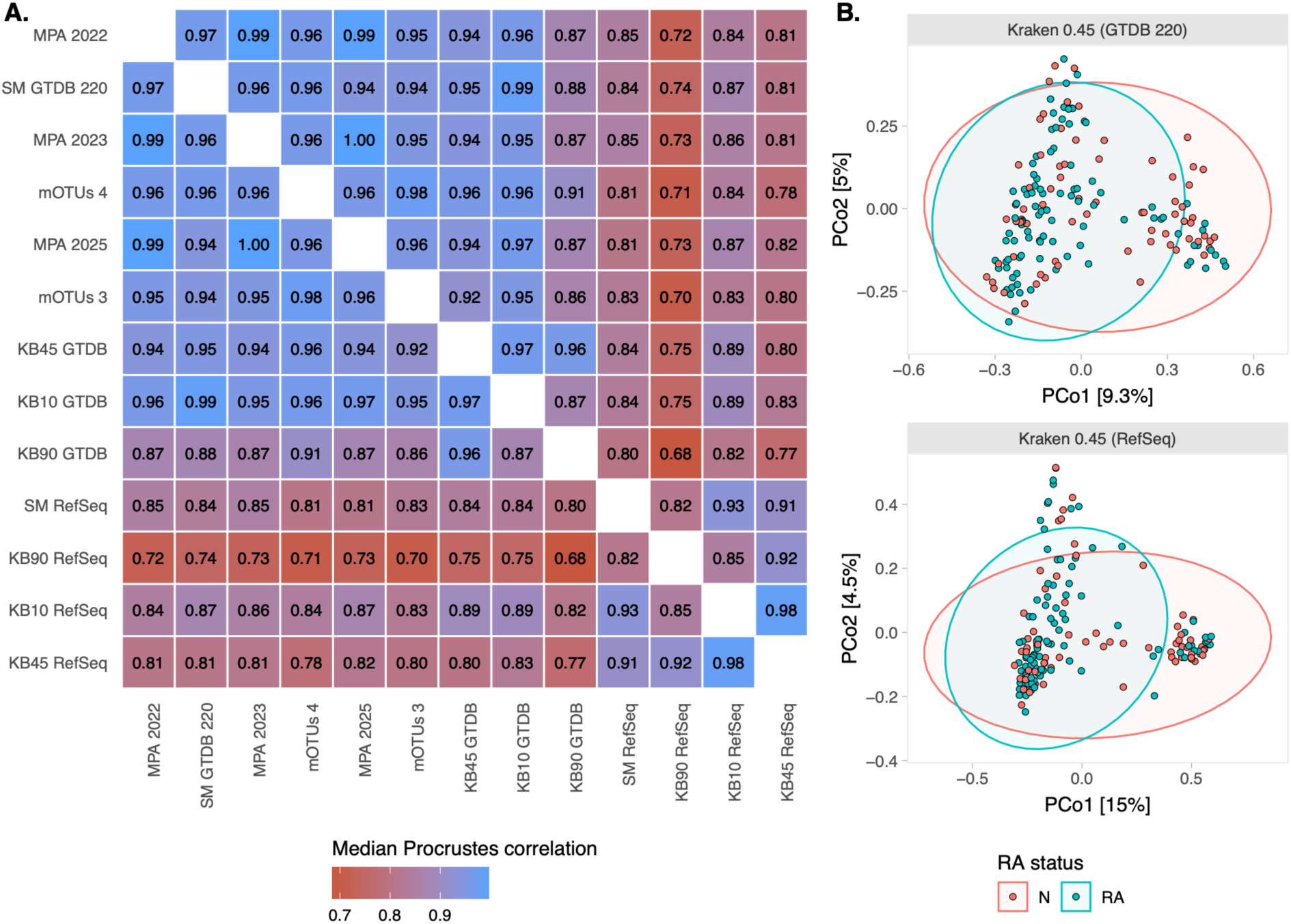
(A) Median Procrustes correlations between pairs of methodologies across eight datasets. Procrustes were calculated on the first 3 axes of the principal coordinates of respective Bray-Curtis dissimilarity matrices. (B) First two principal coordinates (PCo) of the Gut RA dataset dissimilarity matrix clustered by RA status based on taxonomic profiles estimated using Kraken at 0.45 confidence with GTDB (top panel) or RefSeq (bottom panel). SM: Sourmash; KB: Kraken-Bracken; MPA: MetaPhlAn; RA: Rheumatoid arthritis.

To understand which characteristics of the evaluated methodologies may be driving similarity in beta diversity estimates, we plotted the eigenvectors of each Procrustes correlation matrix (**Figure 3**). Overall, methodologies varying only by their database version clustered tightly. All D2M tools clustered on the first axis, except for the Moss dataset that had very poor identifiability using these methods. Database version usually had more influence with mOTUs than with MetaPhlAn, as evidenced by the larger distance between mOTUs3 and 4 on eigenvector 2 for most datasets. D2D tools using RefSeq as a reference databases were the clear outliers, consistently apart from all other methodologies, compared to GTDB and D2M methodologies. One clear exception was Kraken GTDB 0.90, typically isolated from similar methodologies on at least one eigenvector.

**Figure 3.**
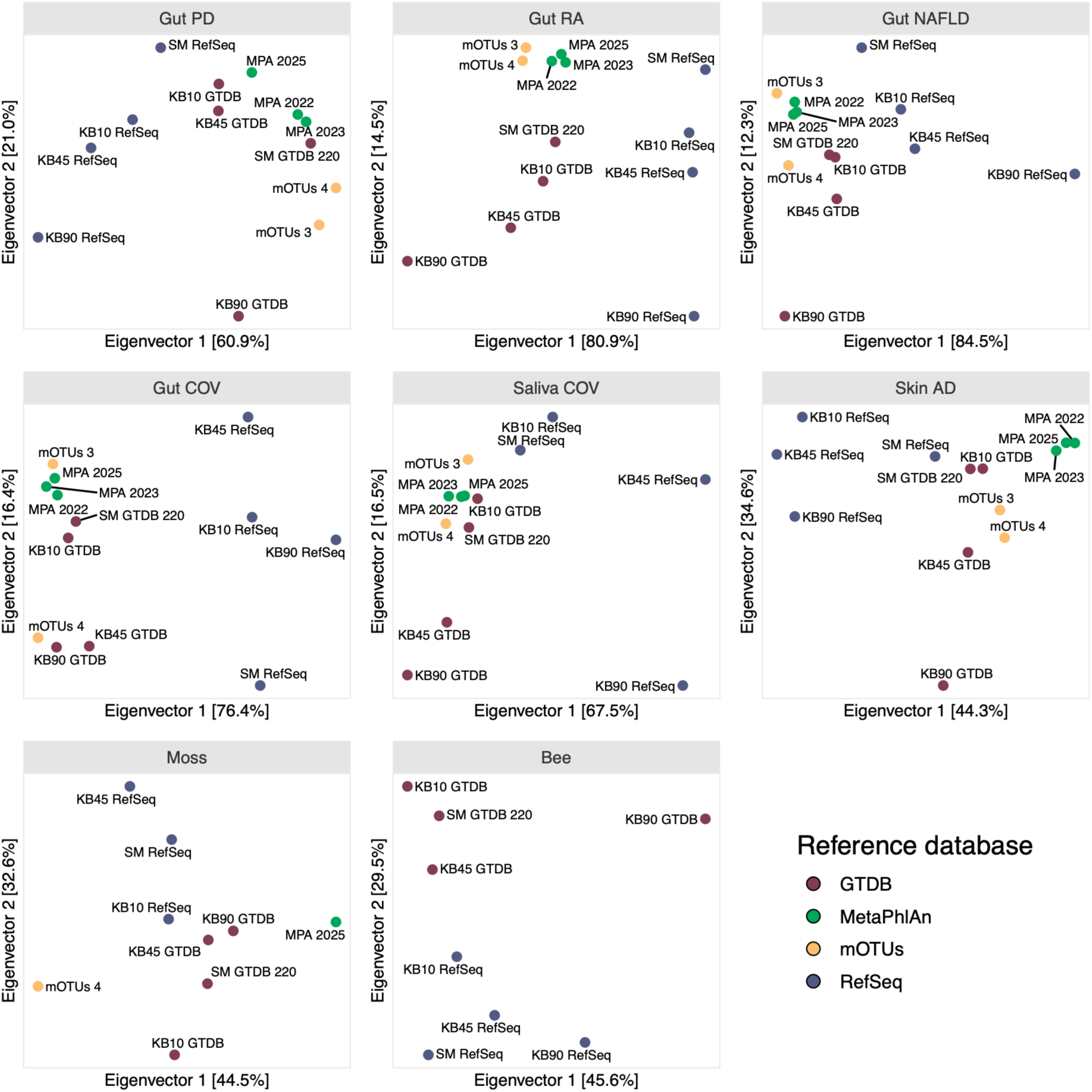
Ordination of Procrustes correlation matrices by dataset.

### Methodology-dependence statistical test interpretation

Comparing Shannon diversity between groups using Wilcoxon rank-sum tests revealed variable effect sizes and p-values across methodologies for most datasets (**Figure 4**). Only the Skin AD and Gut RA datasets showed consistent non-significant results across methodologies. In the Gut COV dataset, difference between groups was only significant (p < 0.01) when using D2M tools or Sourmash GTDB, and effect sizes (rank-biserial correlations) varied from 0.26 to 0.57. In the Gut PD dataset, which wielded higher statistical power with 720 samples, the test was significant for 9 out of 13 methodologies, and the effect sizes ranged from 0.04 to 0.13. With the Saliva COV dataset, only MetaPhlAn led to statistically significant differences between groups (p<0.05), while effect sizes ranged from 0.08 to 0.46. With the Gut NAFLD dataset, effect sizes varied widely across methodologies (0.006 to 0.39), but only Kraken RefSeq 0.90 led to a significant difference between groups (p=0.022). The Moss dataset showed consensus across methodologies, except for mOTUs4, but showed large variations in effect size (0.364 to 0.749, all p<0.001). Similar patterns of discordance in effect sizes and p-values occurred using other diversity indices (**Figure S6**-**Figure S8**). Overall, alpha diversity analysis can be driven by methodological choices, in terms of both significance and effect size.

**Figure 4.**
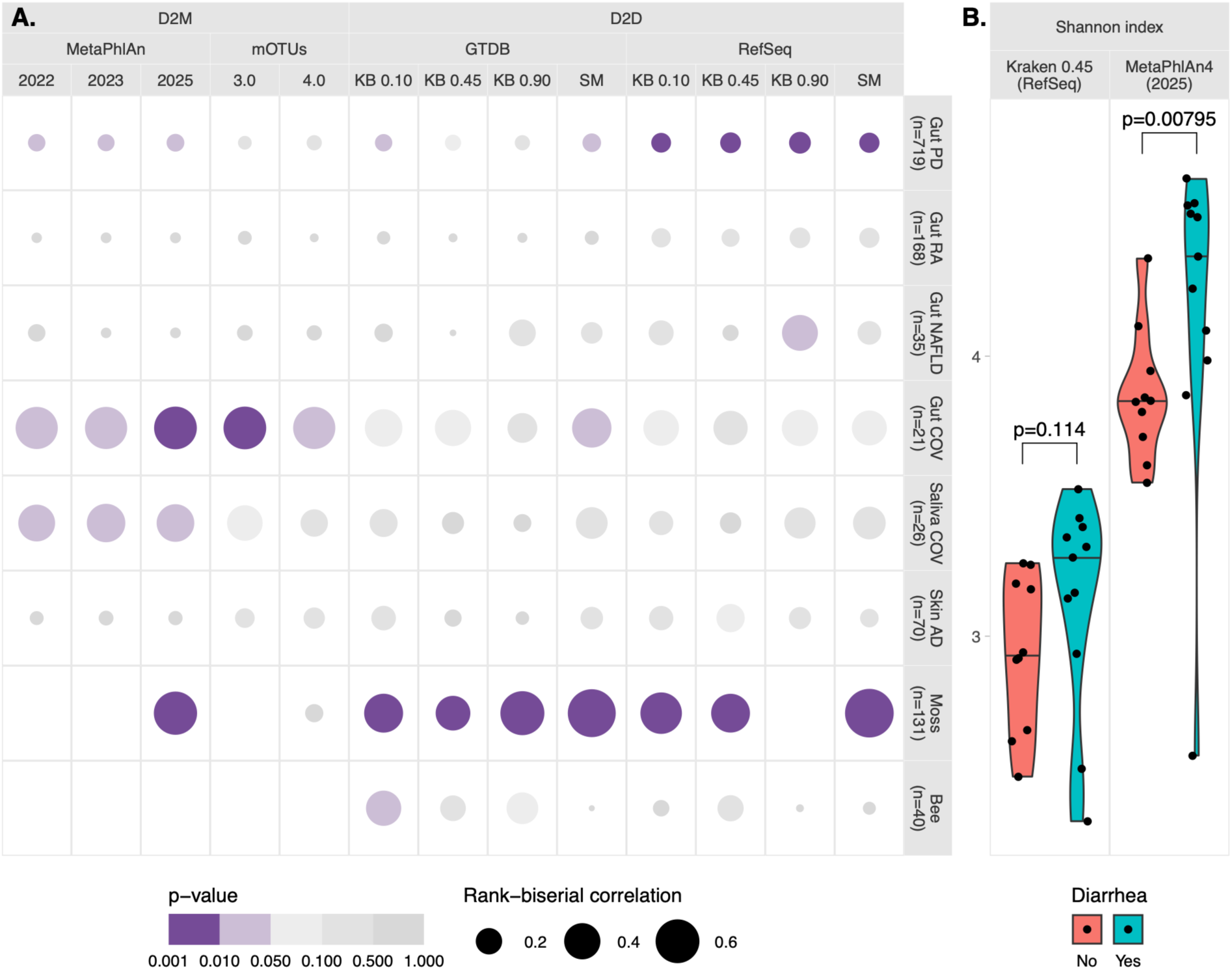
Wilcoxon rank-sum tests comparing Shannon diversity distributions between two groups of samples. A) p-value and rank-biserial correlations by dataset-methodology combination. Empty squares indicate the methodology could not identify any metagenome. B) Distribution of Shannon indices between reported diarrhea groups using KB 0.45 (left panel) or MPA 2025 (right panel). SM: Sourmash; KB: Kraken-Bracken.

To evaluate the sensitivity of beta diversity analysis to methodological choice, we carried out a PERMANOVA on Bray-Curtis dissimilarities for each dataset-methodology combination using the model *Dissimilarity Matrix ∼ Group* (**Figure 5**). The most striking differences across methodologies were the range of R^2^ (explained variance) observed within individual datasets. Despite these differences, most methodologies agreed whether the grouping variable was significantly associated with inter-sample dissimilarities. The Gut COV dataset was the least consistent across methodologies, with D2D tools using RefSeq all yielding non-significant results (*p* > 0.05) with lower R^2^ compared to other methodologies. However, no consistent pattern of methodology effects was observed across datasets. In the case of the Gut NAFLD dataset, all methodologies agreed on statistical significance (p < 0.001). Yet, using D2D tools with RefSeq produced R^2^ between 0.130 and 0.144, whereas D2M tools yielded much smaller values, between 0.072 and 0.080.

**Figure 5.**
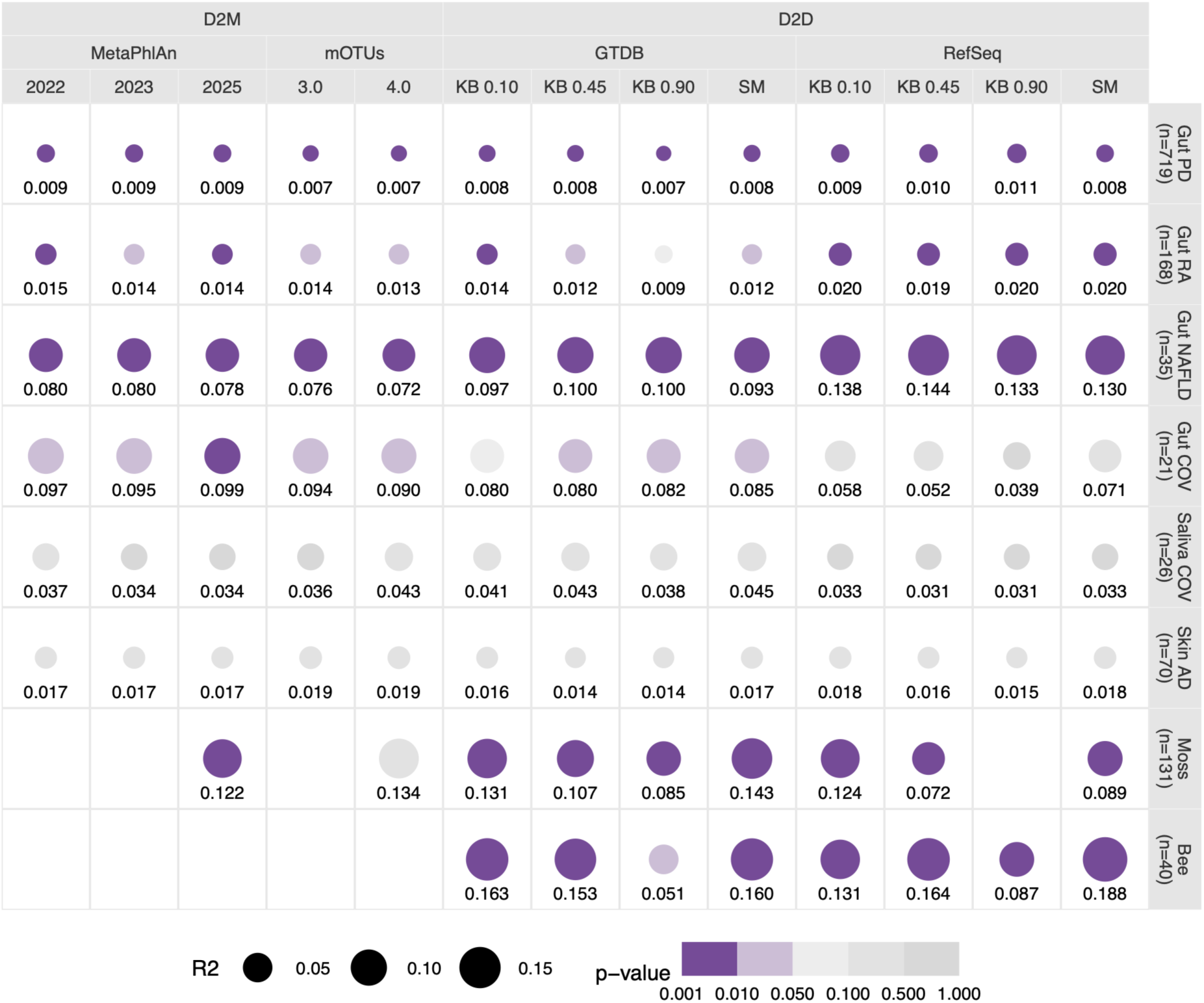
Results of PERMANOVA tests on multivariate regression of the Bray-Curtis dissimilarity matrix on a binary variable representing the predefined groups of samples within each dataset-methodology combination. *SM: Sourmash; KB: Kraken-Bracken*

Overall, methodological choices substantially influenced alpha and beta diversity indices, though the impact on statistical tests using a two-group design was much more pronounced for alpha diversity.

## DISCUSSION

In this work, we address a literature gap by describing four taxonomic profilers for shotgun metagenomics and examining how their characteristics can co-vary with diversity assessment of microbiomes. Applying these tools to 1,221 metagenomes spanning eight datasets, we show how methodological choices affect alpha and beta diversity measurements and their downstream statistical outcomes. We show that study conclusions can shift depending on which taxonomic profiler is chosen and how it is applied, particularly for alpha diversity. Unlike benchmark studies relying on simulated metagenomes, this work focusses on real metagenomes, reflecting the full complexity of microbiomes rather than the current, incomplete knowledge of them. By doing so, we complement the valuable benchmarking question “which tool is the most performant?” with the question “how do methodological choices affect what we conclude?”, providing a relatable angle to researchers hoping to apply these tools to real data. While metagenomics continues to grow in popularity and accessibility, published resources guiding researchers through the analytical landscape are increasingly needed, as is awareness of the importance of methodological design and a better understanding of bioinformatic tools. We hope this work equips researchers with the means to avoid inadequate analyses, cherry-picking, poor reproducibility, and misleading conclusions.

When measuring alpha diversity, regardless of the reference database used, the choice of taxonomic profiler introduces substantial variations in diversity estimation. When controlling for reference database size and sensitivity parameter trade-offs, no tool consistently over- or under-estimated diversity relative to others. Shannon and Inverse Simpson indices were more sensitive to the choice of taxonomic profiler than to the choice of reference database, particularly when comparing Kraken and Sourmash. In contrast, richness and Tail indices were especially tool-sensitive when RefSeq was used as a reference. This suggests that alpha diversity co-varies with the algorithmic strategy (choice of tool) and, when species count carries relatively more weight (i.e., with richness and Tail), with the breadth of the reference database (or, indirectly, sample identifiability). Importantly, reference databases cannot be interchanged between D2M tools, preventing us from de-confounding their influence. It is therefore possible that the strategy used for including unknown species (i.e., meta-OTUs for mOTUs vs. SGBs for MetaPhlAn) is driving some of the observed variability. Interestingly, tool sensitivity in D2M taxonomic profilers was significantly lower, despite the mOTUs4 reference database containing over two times as many species as MetaPhlAn’s 2025 database version (**Table 3**).

The alpha diversity indices themselves varied substantially between methodologies. The outcomes of hypothesis testing based on these indices varied too, though much less with beta diversity. More surprisingly, we found that even tools designed to estimate the same type of relative abundance, using the same reference database, can produce diversity estimates diverging sufficiently to lead to different statistical conclusions. However, variability in measured diversity could reflect dataset characteristics such as biome of origin, phylogenetic diversity, or even sequencing depth, though quantifying it would require a larger number of independent datasets. We identify this as potential for future work. These patterns of sensitivity to methodological choices underscore the need for exercising caution when comparing alpha diversity across studies that use different methodologies. Without transparent reporting of methodology parameters, comparability of scientific conclusions based on the analysis of alpha diversity is limited.

Beta diversity told a different story: although the measured indices themselves were sensitive to methodological choices (**Figure 3**), this sensitivity rarely affected statistical outcomes with the datasets used here (**Figure 5**). For D2M methodologies, pairwise sample dissimilarities were nearly insensitive to the choice of tool or database version, based on Procrustes analyses. This suggests that these tools tend to produce similar amounts of inter-sample variability. D2D methodologies were less stable, with ordinations driven by reference database identity, sensitivity parameters, and possibly algorithmic strategy. Despite these approach-specific observations, PERMANOVA significance and effect size remained stable across almost all methodologies (**Figure 5**). However, this apparent stability does not constitute evidence that PERMANOVA outcomes are robust to methodology; only that observable changes in ordination space do not necessarily result in methodology-dependent conclusions. Moreover, the Kraken-Sourmash comparison suggests that more comprehensive databases, such as GTDB, may improve the resolution of beta diversity analyses by capturing a broader range of distinguishing features across microbial communities. However, the impact of database completeness warrants more thorough investigation, particularly using unpublished metagenomic datasets that reference databases will not have been trained on.

## CONCLUSION

Taken together, these results underscore the substantial influence of methodological choices on the assessment of microbial taxonomic community composition and diversity with short-read metagenomes. Consistent with previous findings [27,39,71], our results highlight that the reference database is a major driver of this influence. Larger reference databases may reduce sensitivity to the choice tool when the D2D approach is chosen, a promising trend given the rapid expansion of available genomic resources [72]. Furthermore, novel tools such as Sylph [73], which estimates both taxonomic and sequence abundances from a single, modifiable database, may reveal additional sources of methodological variability worth characterising in future work.

The increasing diversity of bioinformatic tools and the growing accessibility of running multiple *in silico* experiments on the same samples raises the risk of inadvertent [74] or deliberate cherry-picking [75]. In a field where outcomes are demonstrably sensitive to methodological choices, transparent reporting and justification of methodological decisions are essential for reproducibility and interpretability. In light of our results, we recommend that researchers: (1) review the available tools and current benchmarks to choose appropriately given their research objectives; (2) state and justify the choice of tool(s), parameters, and reference database(s) in their methods section; (3) assess the robustness of their conclusions to choice of reference database; and (4) exercise caution when interpreting and comparing diversity across studies, knowing that the observed signals may be influenced by methodological choices.

Future work incorporating more datasets and a broader range of methodologies could study how methodological parameters interact with dataset characteristics such as biome of origin, taxonomic diversity, and sequencing depth, enabling more targeted recommendations. Valuable research directions include evaluating the sensitivity of complementary diversity indices (e.g., Faith [76] for alpha diversity; UniFrac [63] and Aitchison distances [64] for beta diversity) to methodological variation, investigating the impact of methodology on differential abundance analyses, and assessing how incorporating MAGs from a study’s own metagenomes into the reference database affects conclusions. Ultimately, we hope this work empowers researchers to make more informed, transparent, and reproducible choices in metagenomic analyses, ensuring that biological interpretations are driven by data, not by tools.

## ACKNOWLEDGEMENTS

JRL was supported by a Natural Sciences and Engineering Research Council of Canada (NSERC) grant, a Fonds de Recherche du Québec – Nature et Technologies (FRQNT) grant, and a Graduate Research Excellence Scholarship Program (Fondation Université de Sherbrooke) scholarship. PÉJ holds a Fonds de Recherche du Québec – Santé (FRQS) Research Scholar Career Award. ILL is supported by a Canadian Research Chair T2 (CRC-2023-00177).

We thank Jean-François Lucier (Département de Biologie, Faculté des Sciences, Université de Sherbrooke) for his support with software implementation on high-performance computers. We also thank Jean-Charles Pasquier and his team for the opportunity to work with the PROVID datasets (herein Gut COV and Saliva COV), which had not been released. We thank Sébastien Rodrigue (Département de Biologie, Faculté des Sciences, Université de Sherbrooke) for his involvement and discussions contributing to this project.

## AUTHOR CONTRIBUTIONS

JRL and ILL conceived the study. JRL performed the bioinformatic evaluations and wrote all the scripts. FGB, ILL and PÉJ contributed to the analyses. All authors revised and agreed with the contents herein.

## AVAILABILITY OF DATA AND MATERIALS

The datasets supporting the conclusions of this article are available under the following BioProject accessions PRJNA834801 (Gut PD), PRJNA328258 (Gut NAFLD), PRJNA854811 (Skin AD), PRJEB112409 (Saliva COV and Gut Cov), PRJNA356102 (Gut RA), PRJEB76464 (Moss) and PRJNA685398 (Bee), at the *European Nucleotide Archive*, https://www.ebi.ac.uk/ena. All code written for this study is available at the following Github reposiory: https://github.com/jorondo1/MethodsComparison.

## SUPPORTING INFORMATION

**Figure S1.** Median absolute deviations in Procrustes correlations between pairs of methodologies across eight datasets.

**Figure S2.** Distribution of Inverse Simpson index in samples across pairs of methodologies.

**Figure S3.** Distribution of the Tail statistic in samples across pairs of methodologies.

**Figure S4.** Distribution of richness in samples across pairs of methodologies.

**Figure S5.** Distribution of differences in scaled diversity indices between pairs of methodologies.

**Figure S6.** Results of Wilcoxon tests comparing the distribution of richness between two predefined groups of samples within each dataset-methodology combination.

**Figure S7.** Results of Wilcoxon tests comparing the distribution of the Inverse Simpson index between two predefined groups of samples within each dataset-methodology.

**Figure S8.** Results of Wilcoxon tests comparing the distribution of the Tail index between two predefined groups of samples within each dataset-methodology combination.

**Table S1.** Pitman-Morgan tests comparing the variances of the distribution of differences in scaled diversity indices between pairs of methodologies.

