## Supplementary tables and figures for "Taxonomic profilers and their influence on metagenomic diversity analyses"

### SUPPLEMENTARY FIGURES

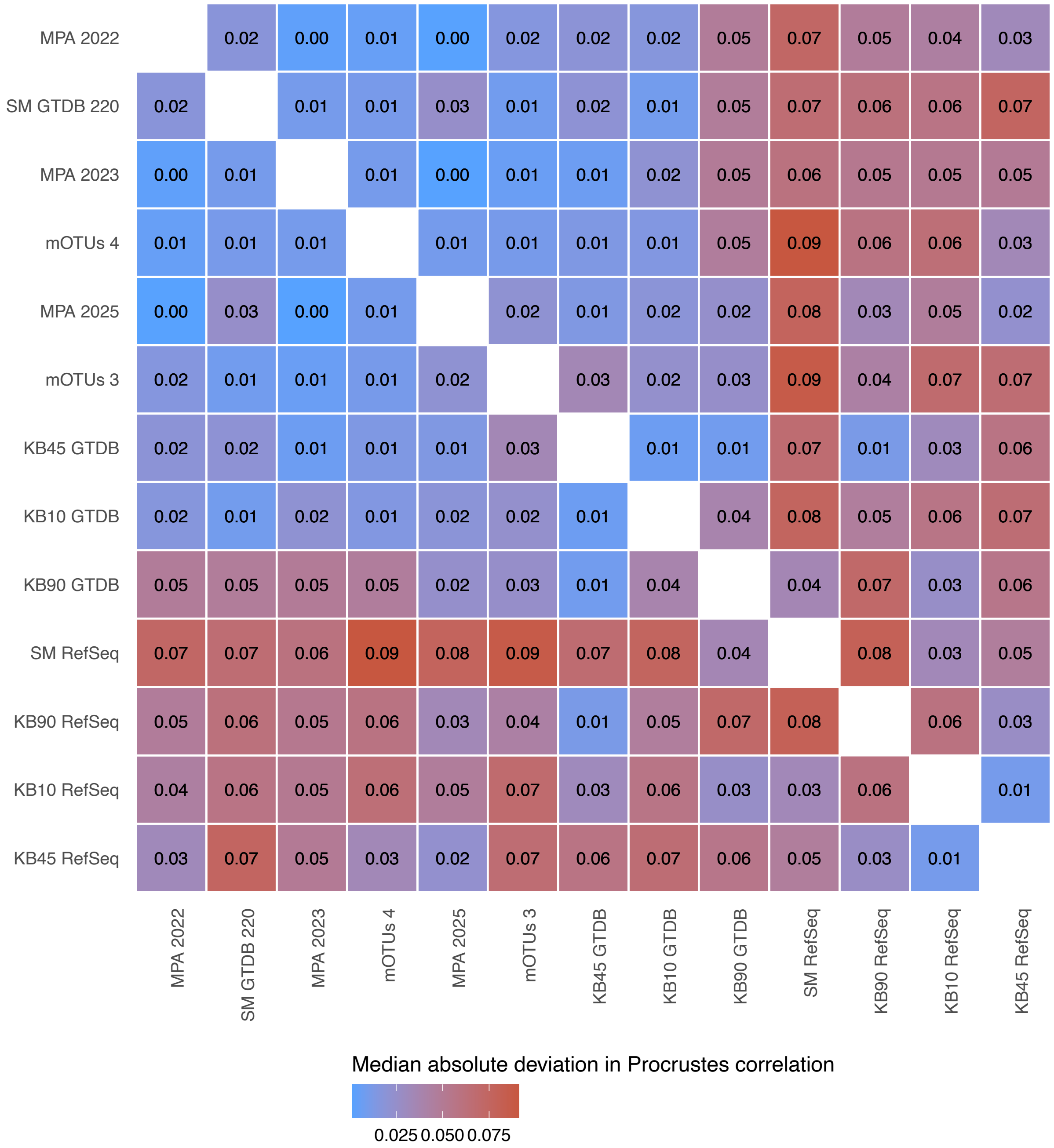


**Figure S1.** Median absolute deviations in Procrustes correlations between pairs of methodologies across eight datasets. Procrustes were calculated on the first 3 axes of the principal coordinates of respective Bray-Curtis dissimilarity matrices.

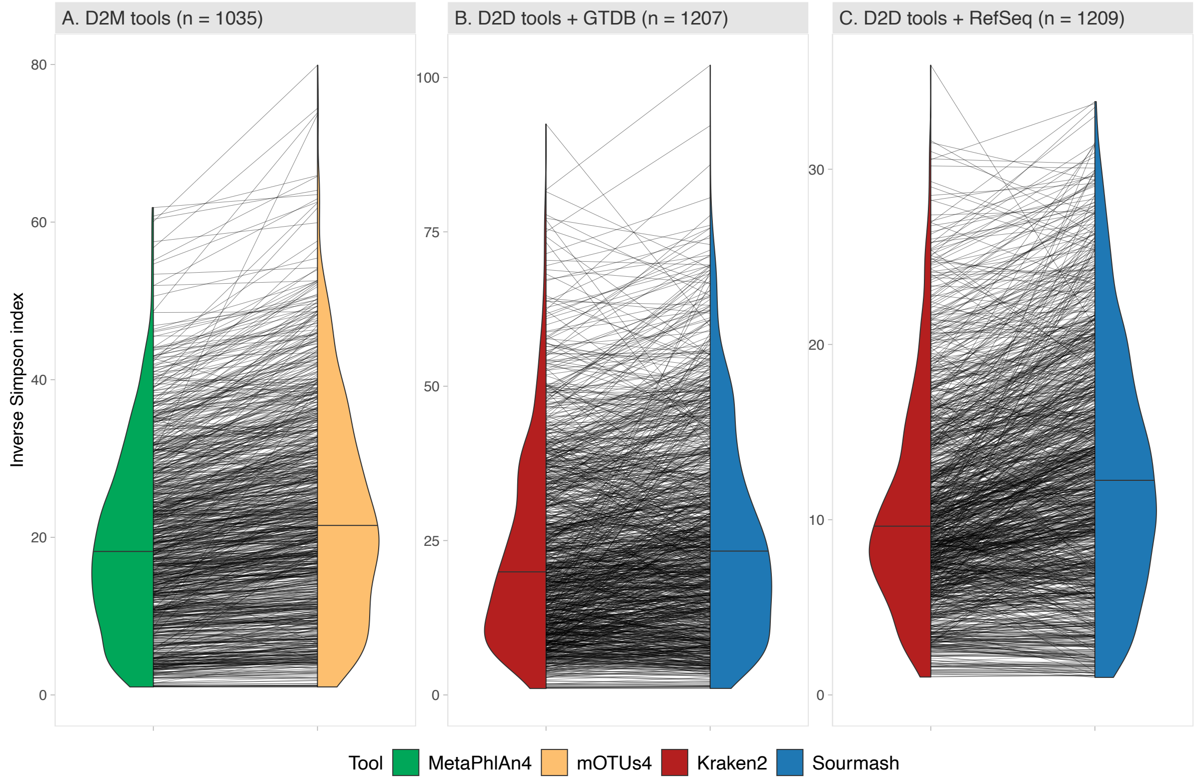


**Figure S2.** Distribution of Inverse Simpson index in samples across pairs of methodologies. Kraken was set at a confidence threshold of 0.45.

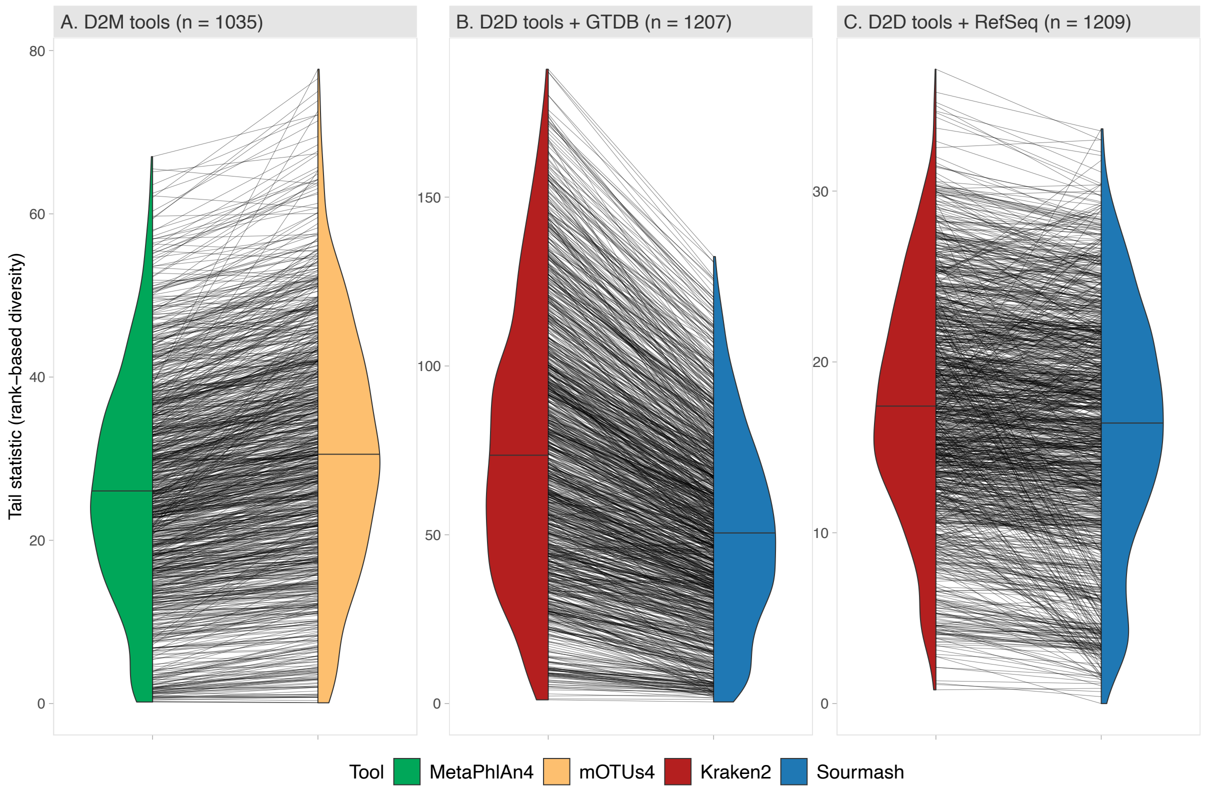


**Figure S3.** Distribution of the Tail statistic in samples across pairs of methodologies. Kraken was set at a confidence threshold of 0.45.

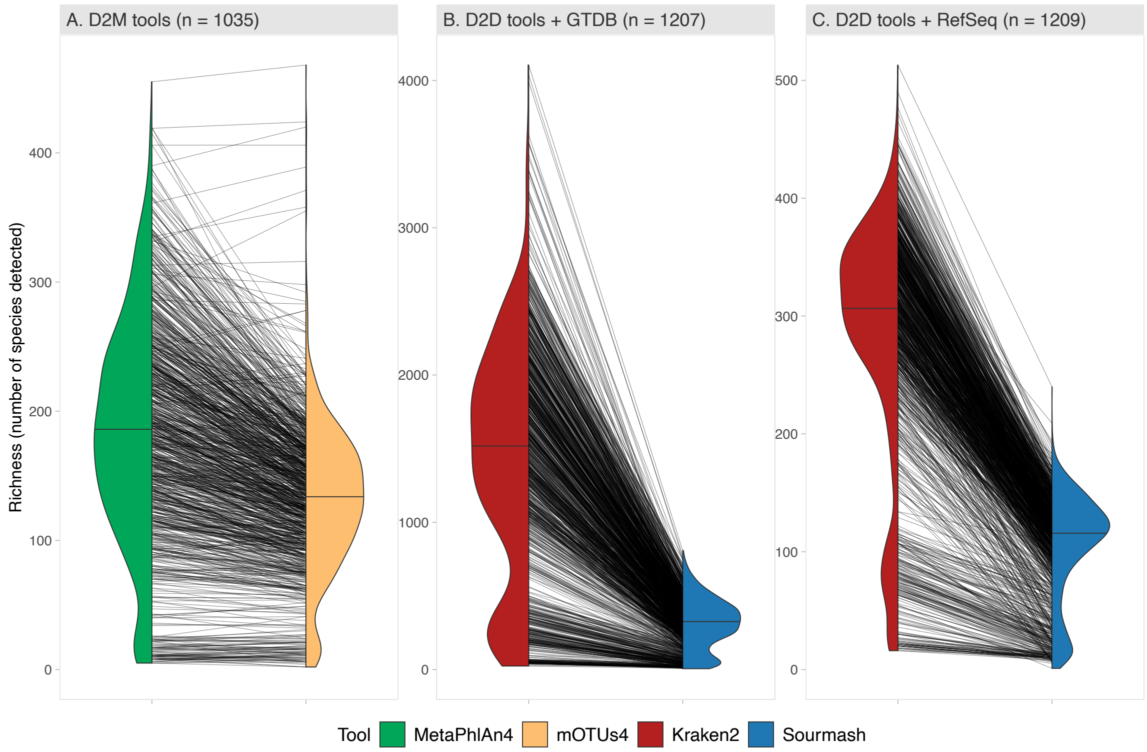


**Figure S4.** Distribution of richness in samples across pairs of methodologies. Kraken was set at a confidence threshold of 0.45.

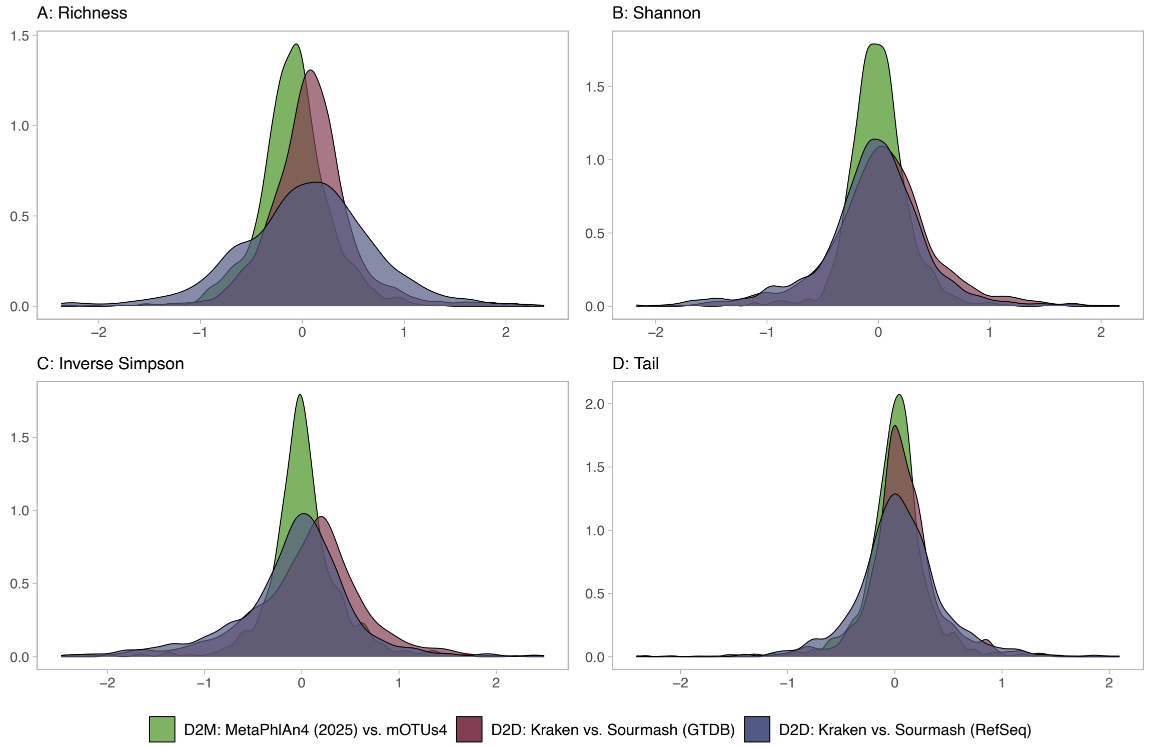


**Figure S5.** Distribution of differences in scaled diversity indices between pairs of methodologies.

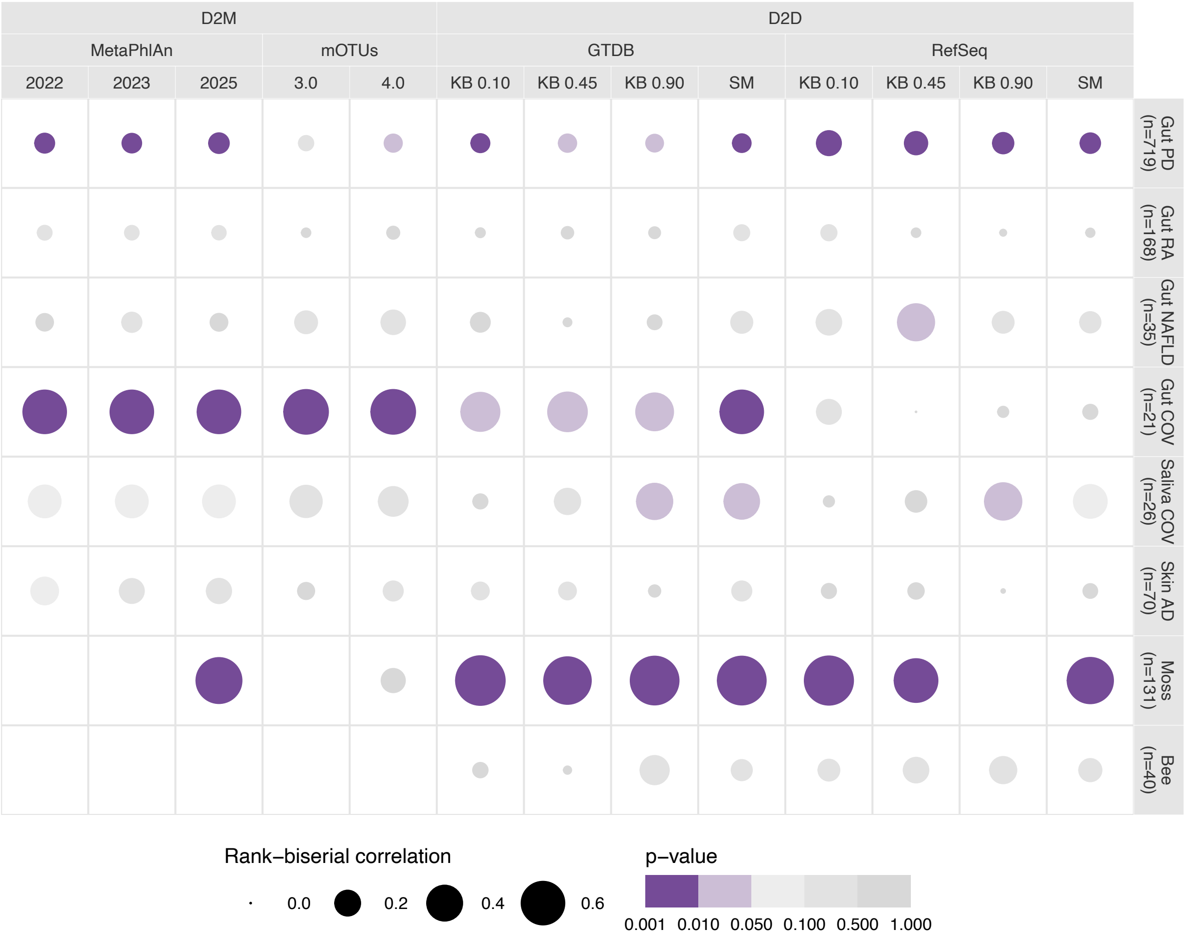


**Figure S6.** Results of Wilcoxon tests comparing the distribution of richness between two predefined groups of samples within each dataset-methodology combination. *SM: Sourmash; KB: Kraken-Bracken.*

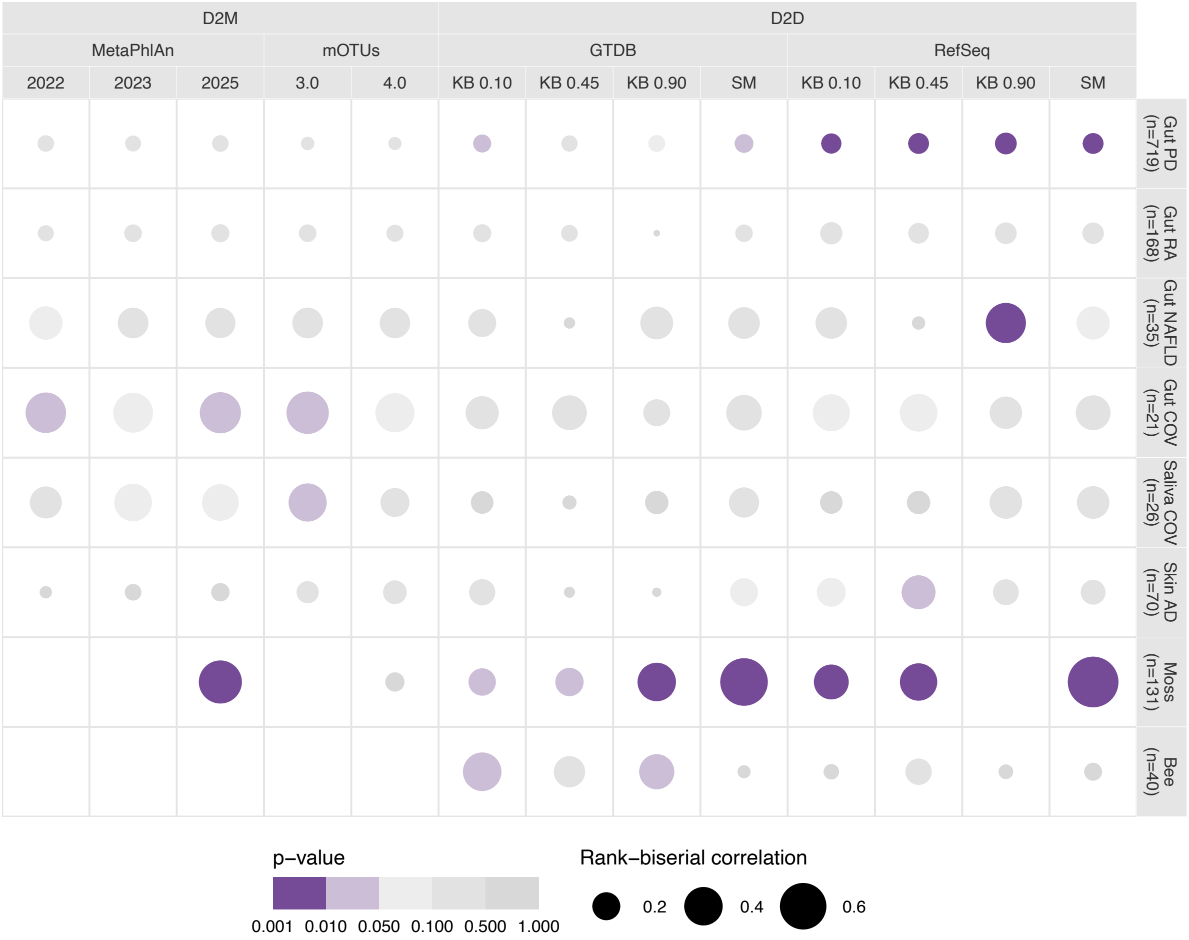


**Figure S7.** Results of Wilcoxon tests comparing the distribution of the Inverse Simpson index between two predefined groups of samples within each dataset-methodology combination. *SM : Sourmash; KB : Kraken-Bracken; MPA : MetaPhlAn.*

13

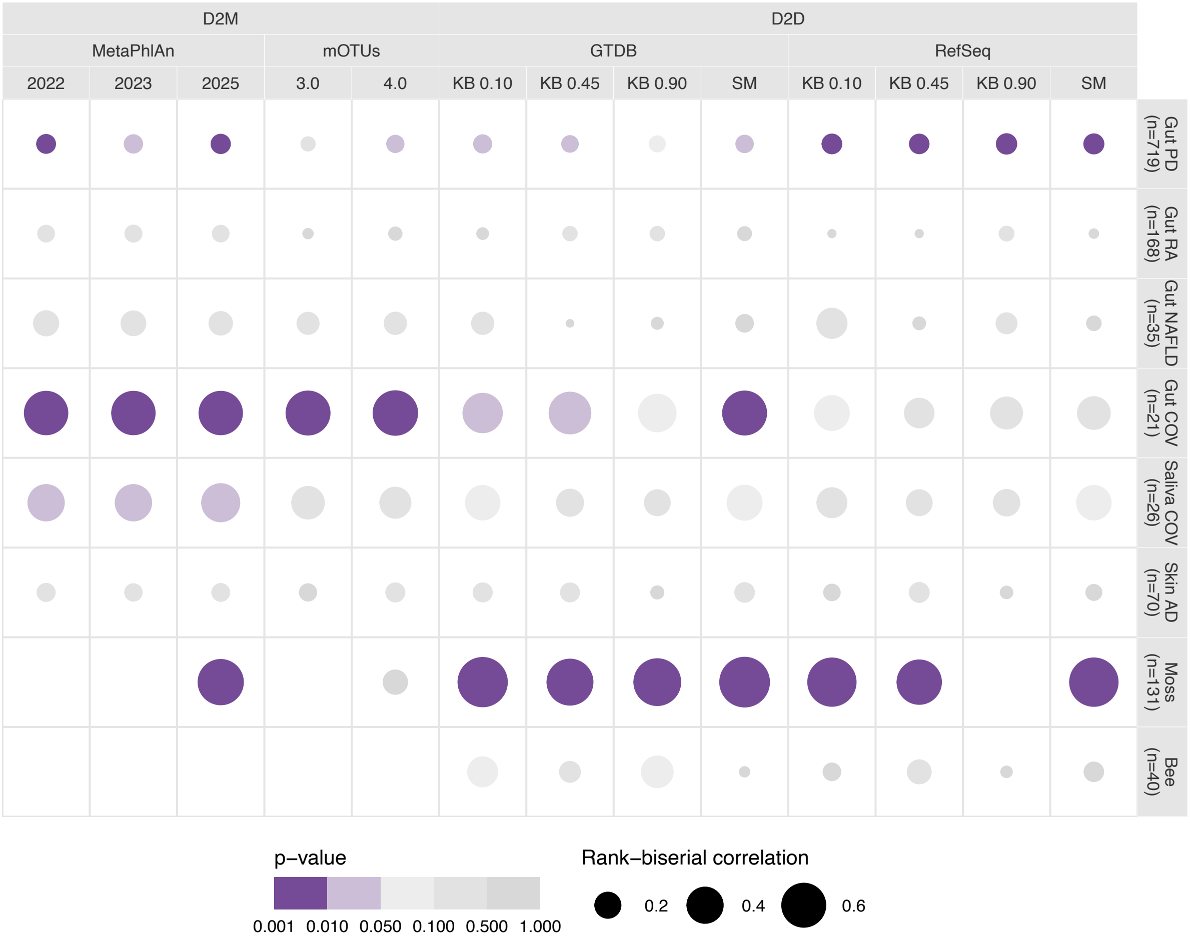


**Figure S8.** Results of Wilcoxon tests comparing the distribution of the Tail index between two predefined groups of samples within each dataset-methodology combination. *SM : Sourmash; KB : Kraken-Bracken; MPA : MetaPhlAn.*

*SUPPLEMENTARY TABLES*

**Table S1.** Pitman-Morgan tests comparing the variances of the distribution of differences in scaled diversity indices between pairs of methodologies.

| | | | Variance | | | |
|---|---|---|---|---|---|---|
| | **Method pair 1** | **Method pair 2** | **Pair 1** | **Pair 2** | **Difference** | **Adj. p[2]** |
| **Richness** | MetaPhlAn4[1] vs. mOTUs4 | Kraken vs. Sourmash (GTDB) | 0.148 | 0.191 | 0.043 | 0.0000 |
| | MetaPhlAn4 vs. mOTUs4 | Kraken vs. Sourmash (RefSeq) | 0.148 | 0.453 | 0.305 | 0.0000 |
| | Kraken vs. Sourmash (GTDB) | Kraken vs. Sourmash (RefSeq) | 0.240 | 0.475 | 0.235 | 0.0000 |
| **Shannon** | MetaPhlAn4 vs. mOTUs4 | Kraken vs. Sourmash (GTDB) | 0.080 | 0.174 | 0.094 | 0.0000 |
| | MetaPhlAn4 vs. mOTUs4 | Kraken vs. Sourmash (RefSeq) | 0.080 | 0.235 | 0.155 | 0.0000 |
| | Kraken vs. Sourmash (GTDB) | Kraken vs. Sourmash (RefSeq) | 0.258 | 0.275 | 0.017 | 0.4681 |
| **Inverse** | MetaPhlAn4 vs. mOTUs4 | Kraken vs. Sourmash (GTDB) | 0.179 | 0.341 | 0.162 | 0.0000 |
| | MetaPhlAn4 vs. mOTUs4 | Kraken vs. Sourmash (RefSeq) | 0.179 | 0.378 | 0.199 | 0.0000 |
| | Kraken vs. Sourmash (GTDB) | Kraken vs. Sourmash (RefSeq) | 0.491 | 0.496 | 0.005 | 0.8657 |
| **Tail** | MetaPhlAn4 vs. mOTUs4 | Kraken vs. Sourmash (GTDB) | 0.084 | 0.094 | 0.010 | 0.1638 |
| | MetaPhlAn4 vs. mOTUs4 | Kraken vs. Sourmash (RefSeq) | 0.084 | 0.140 | 0.056 | 0.0000 |
| | Kraken vs. Sourmash (GTDB) | Kraken vs. Sourmash (RefSeq) | 0.150 | 0.185 | 0.034 | 0.0001 |

[1]MetaPhlAn used with the 2025 database.

[2]P-values were adjusted using the Holm procedure.
